# Viral dispersal drives changes in population composition

**DOI:** 10.64898/2026.10.12.758528

**Authors:** Katie E. Holmes, Anna Sims, Lucas M. Ferreri, Lucia Ortiz, Baptiste Elie, Katia Koelle, David VanInsberghe, Anice C. Lowen

**Author notes:** Institute of Medical Virology, University of Zurich, Zurich, Switzerland (Baptiste Elie).

## Abstract

In an acute viral infection, a small seeding population expands rapidly over several orders of magnitude, then declines. This vast population growth is accompanied by spatial expansion as infection spreads to new anatomical sites within the host. Evolutionary theory predicts that spatially expanding populations experience more extreme stochastic losses of genetic diversity than spatially unstructured populations. To test this prediction in the context of viral dynamics, we evaluated the impact of spatial expansion on viral population genetic structure and assessed how modulators of viral dispersal affected this structure. To this end, we developed a means of implementing the spatial expansion of influenza A virus in cell culture and applied it to a viral population carrying a diverse genetic barcode. Across this simplified cellular landscape, we found that spatial expansion produces a patchwork of distinct viral subpopulations. We then evaluated the consequences for population structure of two modulators of viral dispersal: extracellular viscosity and a common polymorphism in the viral attachment protein. Both physical constraint and viral phenotype altered the extent of genetic bottlenecking and spatial heterogeneity that developed during expansion, indicating that environmental features and viral traits that modulate dispersal also modulate viral population composition, diversity and evolution.

## Introduction

The evolution of viral populations enables their escape from immune effectors, evasion of antiviral drugs and spread to new host species. While these forms of adaptive evolution are notable for their negative consequences for host populations, they occur against a backdrop of viral evolution driven by stochastic, non-selective processes. These non-selective evolutionary processes play a prominent role in shaping viral populations within-hosts (1, 2), during transmission (3–8) and at the host population level (9–11). This is important because populations strongly impacted by stochastic processes are thought to adapt more slowly, as chance events can lead to the stochastic spread of deleterious mutations and stochastic loss of beneficial mutations. Understanding the relative roles of stochasticity and natural selection in shaping the genetic composition of viral populations is thus critical for anticipating the ability of viral populations to adapt to new and changing environments.

Among stochastic processes, the potency with which genetic drift acts on a population depends on the population’s size and on the extent to which reproductive output varies across individuals due to chance (12). Another important non-selective factor impacting population evolution is spatial structure, or the non-random arrangement of entities across space. Populations that expand with spatial structure experience higher levels of genetic drift than those proliferating in a spatially unstructured environment, as subpopulations at the leading edge are often very small (13–15). In the context of viruses, a newly infected host can be considered an initially uninhabited range over which a virus population expands. This scenario recurs frequently for acutely infecting viruses, with infection of each new host characterized by a massive expansion from the initial site of infection followed by contraction as the infection is cleared. By this viral analogy to ecological range expansions, the evolutionary dynamics of spatially expanding viral populations should be impacted by the process of genetic drift despite large overall population sizes (16).

Viral dispersal at the within-host level is modulated by the host environment and by viral traits (17–20). Influenza viruses in humans infect the epithelial lining of the respiratory tract and therefore navigate an often-complex topography that differs greatly along the progression from upper to lower respiratory tissues. Influenza virions move within the fluids that line the apical surface of the epithelium, which vary in viscosity through an apical cross-section and across the anatomy of the respiratory tract (21). Their movement over short distances is driven by diffusion and viral attachment to and release from host cells, mediated by the hemagglutinin (HA) and neuraminidase (NA) glycoproteins, respectively (22, 23). Dispersal over long distances is likely facilitated by ciliary action and respiration (24–26). Thus, both conditions of the host environment and viral traits have the potential to modify dispersal distances and the extent of spatial structure. We hypothesize that, through their effects on dispersal, these same factors will also shape population diversity and evolutionary potential.

To investigate the relationships between viral dispersal and population genetic composition, we devised a means of simulating spatial structure in cell culture and then applied this system to examine spatial variation in the composition of a barcoded viral population. Our results reveal that viral dissemination even over a simplified landscape is often associated with losses in population diversity and gives rise to extensive spatial variation in population composition. This effect was sensitive to the viscosity of the extracellular environment and to viral variation in the HA receptor binding domain, revealing the potential for both environmental features and viral phenotypes to modulate the evolutionary potential of dispersing viral populations.

## Results

### Spatial structure of viral populations can be modeled in cell culture

To model spatial structure experimentally, we sought to develop a method in which the location of the source population was well-defined, the ease of dispersal could be controlled, and sink populations at a range of distances from the source could be sampled.

We found that a source population could be placed at the center of a culture dish by seeding infected, adherent cells within a drop of medium and then allowing these cells to attach (**Figure 1A**). Once the source cells had adhered, medium containing uninfected cells could be flooded across the dish to create a monolayer. Inclusion of methylcellulose in this culture medium to increase viscosity offered a means of manipulating the extent of viral dissemination. To allow sampling at explicit locations throughout the dish, glass coverslips could be placed prior to the seeding of cells. At the time of sampling, the coverslips and associated cells could be removed from the dish and processed as desired.

**Figure 1:**
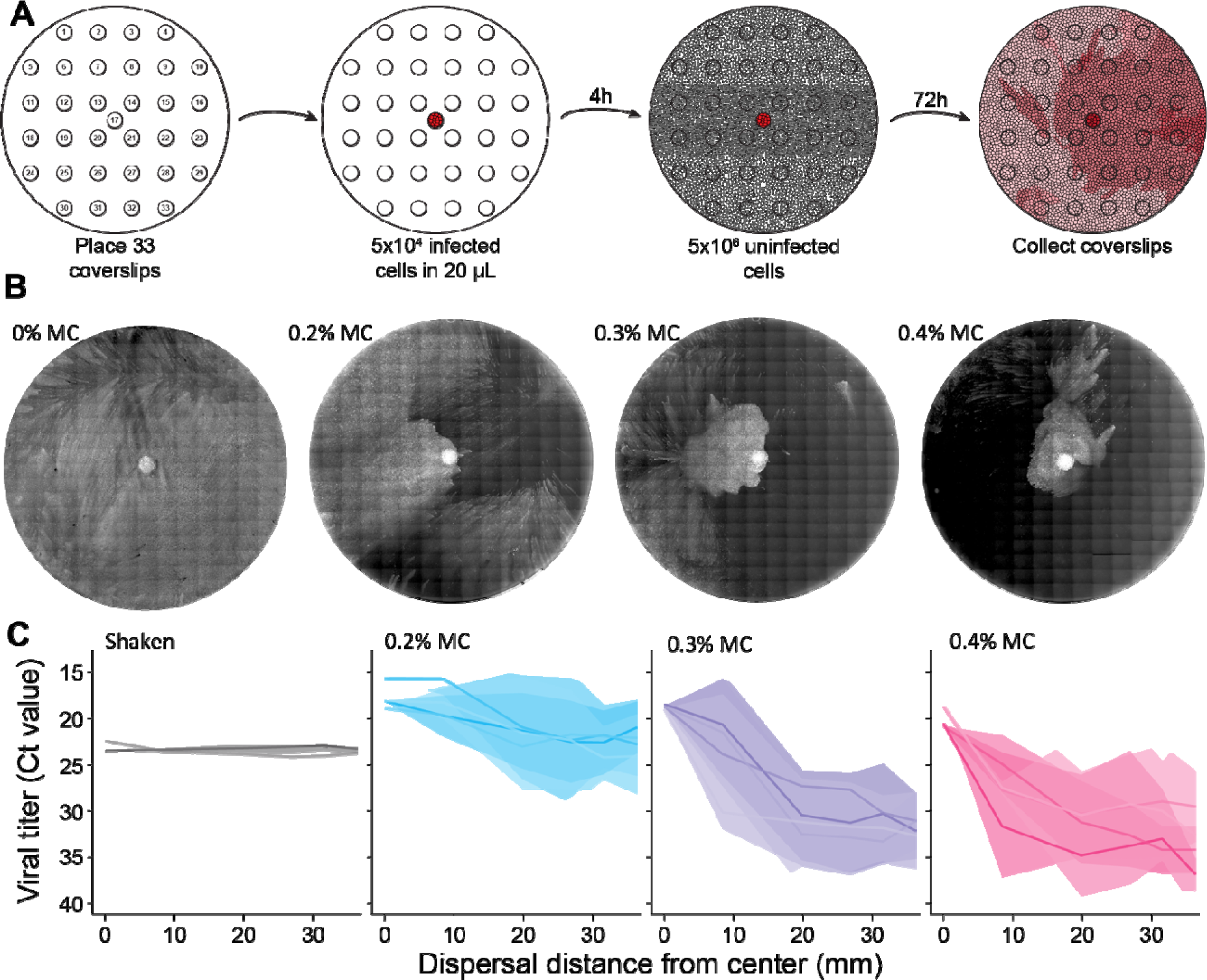
Viral spatial structure develops readily in static culture dishes and can be augmented by increasing viscosity. A) Schematic diagram showing methodology. Small circles represent coverslips; large circles represent 10 cm diameter culture dishes; red coloring shows infected cells. Infected cells are placed in the center (at coverslip no. 17) to seed the source population. B) Images of dispersed viral populations in 10 cm dishes 72 h after the seeding of the central focus. The v/v percentage of methylcellulose in the culture medium is indicated (MC = methylcellulose). C) Relationships between viral load at 72 h post seeding and distance from the central focus for shaken plates with no methylcellulose and static plates at three different methyl cellulose concentrations. Each line represents a single culture dish, with shading showing 95% confidence intervals.

Population density across space was evaluated in two ways. First, to obtain a visual assessment, plates were seeded with a virus encoding a tetra cysteine tag, stained at 72 h post-seeding and then imaged using a tiling approach to capture the full plate (**Figure 1B**). Second, to quantify viral density across space, coverslips were placed prior to viral seeding, collected at 72 h post-seeding and then processed for quantification of viral genomes by RT qPCR (**Figure 1C**). Upon incubation of plates under shaken conditions, virus spread from the source across the full area of the plate. Images of this condition were not obtained because it led to extensive cell death. Conversely, under static conditions, spread was non-uniform and gave rise to substantial heterogeneity in population density (**Figure 1B**). While increasing methylcellulose concentration had little impact on viral load at the source population, it decreased the size of dispersed viral populations (**Figure 1C**).

### Spatially structured spread gives rise to population genetic heterogeneity

To examine how viral propagation over a spatial expanse shapes the genetic composition of the viral population, we used a barcoded virus library as the source population and evaluated barcode composition at 32 peripheral positions across the culture dish.

The barcode library is built into the NA gene segment of influenza A/Panama/2007/99 (H3N2) virus to yield Pan/99 NA-BC (27). This virus population carries abundant synonymous diversity, comprising approximately 4000 distinct genotypes (**Supp Figure 1**). Cells infected at high multiplicity with the Pan/99 NA-BC virus library were used to seed the source population at the center of culture dishes. Dishes were then overlaid with cells in media containing 0.2, 0.3 or 0.4% methylcellulose and then incubated for 72 h to allow for viral dispersal across the cell monolayer. As a comparator, some dishes were agitated continuously throughout the 72 h period to create well-mixed populations. Coverslips were then collected to sample cells at the 32 pre-defined locations on each dish and viral barcodes therein were identified by next generation sequencing.

Whereas viral populations that expanded under agitation were found to be spatially homogeneous in their density (as assessed by Ct values), those that dispersed under static conditions showed variable population density which tended to decline with distance from the source population (**Figure 1C**). Population composition was also homogenous for agitated dishes, with high diversity maintained across the cellular landscape (**Figure 2**). By contrast, static dishes showed marked variation in population composition across space (**Figure 2**). Population composition did not show a clear relationship with distance from the source, likely due to a lack of radial symmetry in dispersal. Instead, one or two sectors of a given dish typically maintained relatively high barcode diversity while the remainder of the area hosted low diversity subpopulations that were highly dissimilar from the source (**Figure 2B**). Despite this local variation, dissimilarity from the source tended to be greater with higher methylcellulose concentration (**Figure 2B**). Modulation of viscosity with methylcellulose also impacted the propensity of a given barcode to be detected across many sampling locations: with higher viscosity, relatively few locations carried a high proportion of barcodes, indicating tighter bottlenecking than seen with lower viscosity (**Figure 2C**). The difference in spatial structure imposed by modulating viscosity is underlined by the more comparable outcomes of low viscosity plates to shaken plates, wherein spatial structure is entirely disrupted (**Figure 2C**). Thus, spatially structured viral dispersal across a simplified, two-dimensional landscape gave rise to strong heterogeneity in population composition, which could be modulated by altering extracellular viscosity.

**Figure 2:**
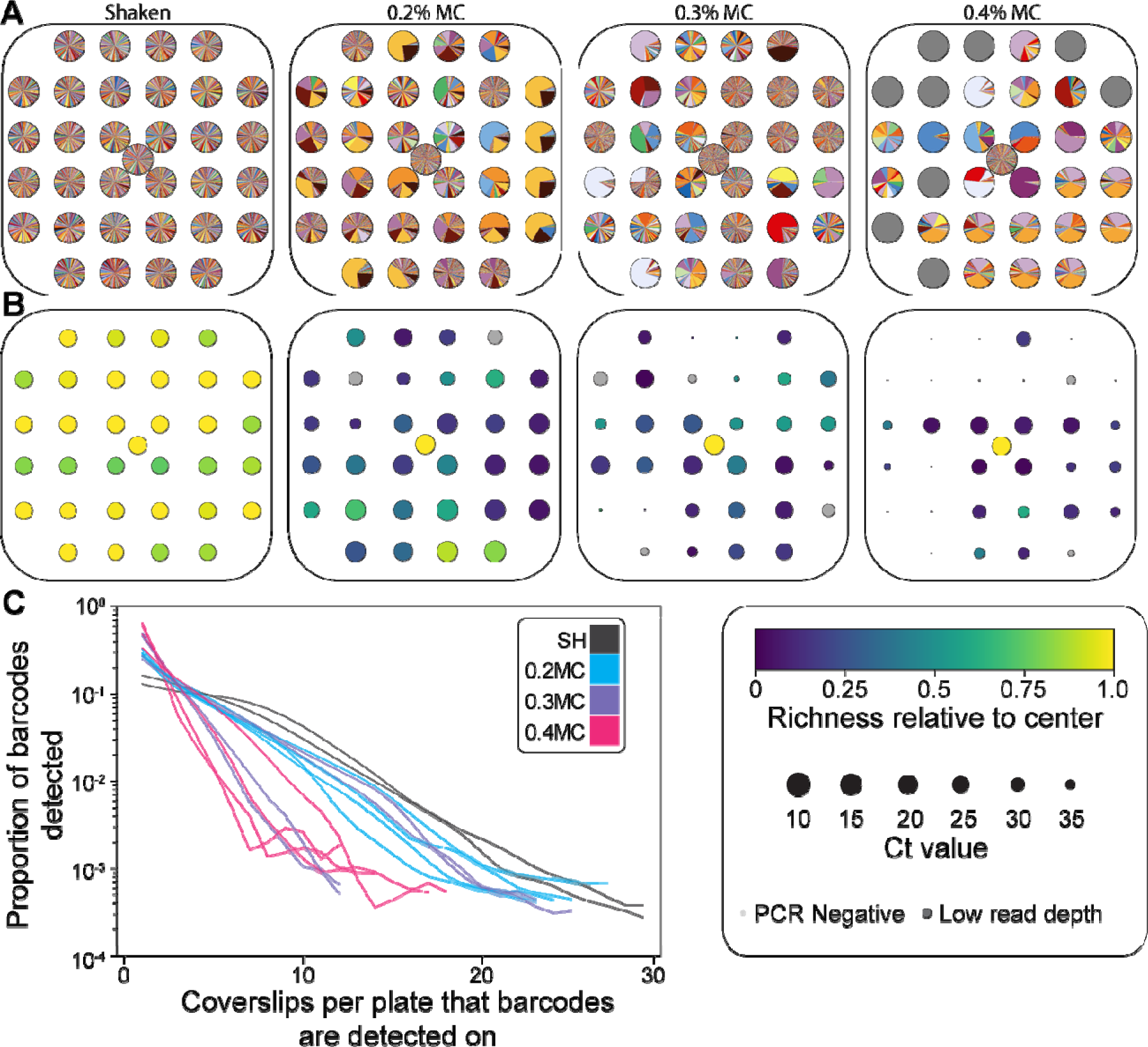
Spatially structured dispersal reduces diversity and precipitates genetic heterogeneity across space. A) Pie charts show barcode composition at 33 locations per cell culture plate. Each color represents a unique barcode and the size of the wedge indicates its frequency within the locally sampled population. Conditions are identified at the top of the panel and representative plates for each condition are shown; all plates are shown in supplementary figure 2. B) Heat maps show barcode richness relative to the center focus, with circles scaled to viral load as measured with Ct values. The same plates are displayed in panels A and B. All plates are shown in supplementary figure 2. C) The proportion of all barcodes that are detected on a given coverslip is plotted against the number of coverslips per plate where that proportion is observed. Each line represents a single cell culture plate, with blue and pink colors representing different methylcellulose (MC) concentrations. Black lines show results from plates incubated with constant shaking (SH), documenting results in the absence of spatial structure. Shading shows the 95% confidence interval of the distribution observed over 1000 sub-samplings of 3000 reads per coverslip.

### Viral phenotypes modulate the relationship between population expansion and population composition

Expansion of influenza virus populations to new locations within a host is mediated by the movement of virions through extracellular space. To initiate a new infection, these virions attach to target cells via the receptor binding domain of the hemagglutinin (HA) protein. Owing to immune pressure that drives its antigenic evolution, this region of the HA protein is highly variable. We assessed the implications of variation in the HA receptor binding domain for viral dispersal and for the changes in population composition that accompany dispersal.

A variant of Pan/99 NA-BC virus was prepared with a change at HA position 159 that is commonly seen during antigenic evolution of H3N2 subtype viruses in humans (HA 159D). Using standard methods of inoculation and culture, Pan/99 NA-BC and Pan/99 NA-BC-HA_159D_ viruses showed comparable amplification from a low multiplicity of infection in MDCK cells (**Figure 3A**). Conversely, in a plaque assay, where spatial structure is imposed with an agar-based overlay, the Pan/99 NA-BC-HA_159D_ variant produced larger plaques (**Figure 3B**). A dissemination phenotype was again apparent when spread from a central focus was evaluated in 10 cm culture dishes, with Pan/99 NA-BC-HA_159D_ accessing a greater expanse than Pan/99 NA-BC virus (**Figure 3C, D**).

**Figure 3:**
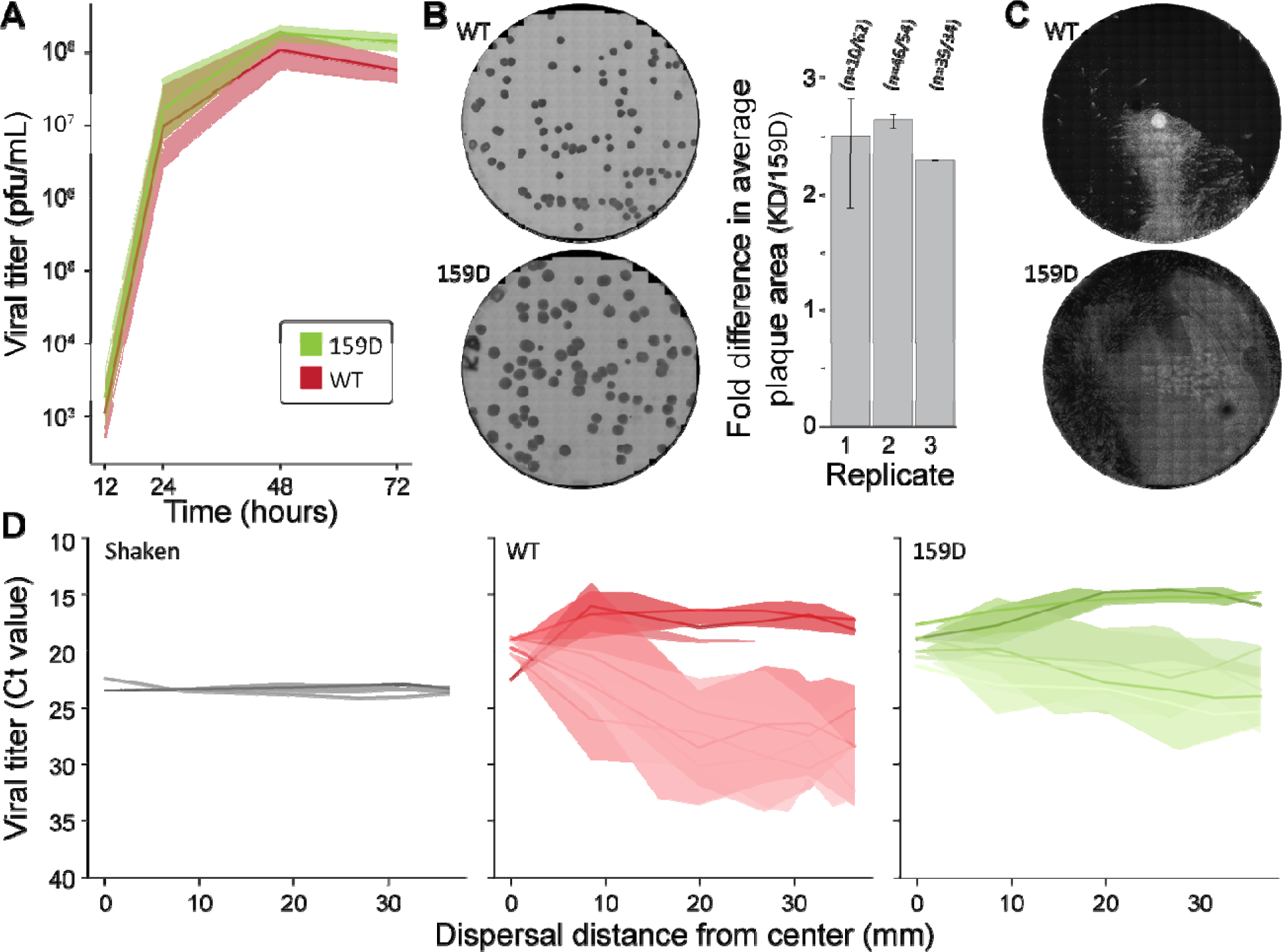
Changes in the HA receptor binding domain modulate viral dispersal. A) Viral replication in MDCK cells following bulk inoculation at MOI=0.01 PFU/cell and incubation under standard (low viscosity) conditions. Three culture dishes per virus were sampled and results are plotted as the mean with 95% confidence intervals. B) Representative images are shown of the plaques formed by WT and 159D variant viruses and the mean fold difference in plaque size is plotted for three replicate plaque assays. The number of plaques measured is indicated above each bar (159D/WT). Error bars show standard deviation. C) Representative images showing the dispersal of WT and 159D variant viruses from a center focus at 72 h post seeding, with 0.2% MC. D) Relationships between viral load at 72 h post seeding and distance from the central focus for the WT virus in the shaken condition (black) and for the WT (red) and 159D (green) variant viruses following 72 h of spatially structured expansion in 0.2% MC. Each line represents a single culture dish, with shading showing 95% confidence intervals. Data for the shaken condition are reproduced from Figure 1 for reference.

The implications of this HA-mediated phenotypic change for the evolution of population composition during expansion was evaluated by implementing the dispersal of Pan/99 NA-BC and Pan/99 NA-BC-HA_159D_ viruses in parallel and then collecting spatially explicit cellular subpopulations for analysis. Both viruses spawned subpopulations with barcode compositions that were often low in diversity and highly dissimilar from the source (**Supplementary Figure 2**). However, individual barcodes within the Pan/99 NA-BC virus population had more limited reach than those in the Pan/99 NA-BC-HA_159D_ population. This was apparent in the increased proportion of barcodes that co-occurred on a given number of coverslips when barcodes were paired with the HA_159D_ variant instead of the HA_WT_ variant (**Figure 4A**). To assess whether differences in population composition between these viral variants could be attributed to close-range or long-distance dispersal, we evaluated the proportion of barcodes shared between two classes of coverslips: those that are nearest to a focal coverslip and those that are furthest. We found that coverslips in both distance classes that were infected by the Pan/99 NA-BC-HA_159D_ virus shared a greater proportion of barcodes than those infected with Pan/99 NA-BC virus (p<0.01, Welch’s t-test) (**Figure 4B**). In addition, for both viral variants, furthest coverslips shared a lower proportion of barcodes compared to nearest coverslips. This effect was more extreme for Pan/99 NA-BC compared to Pan/99 NA-BC-HA_159D_, however (**Figure 4B**). These results suggest that variation in the HA receptor binding domain, like that commonly seen in circulating influenza viruses, can alter not only the potential for dissemination, but also the evolutionary change that accompanies dissemination.

**Figure 4:**
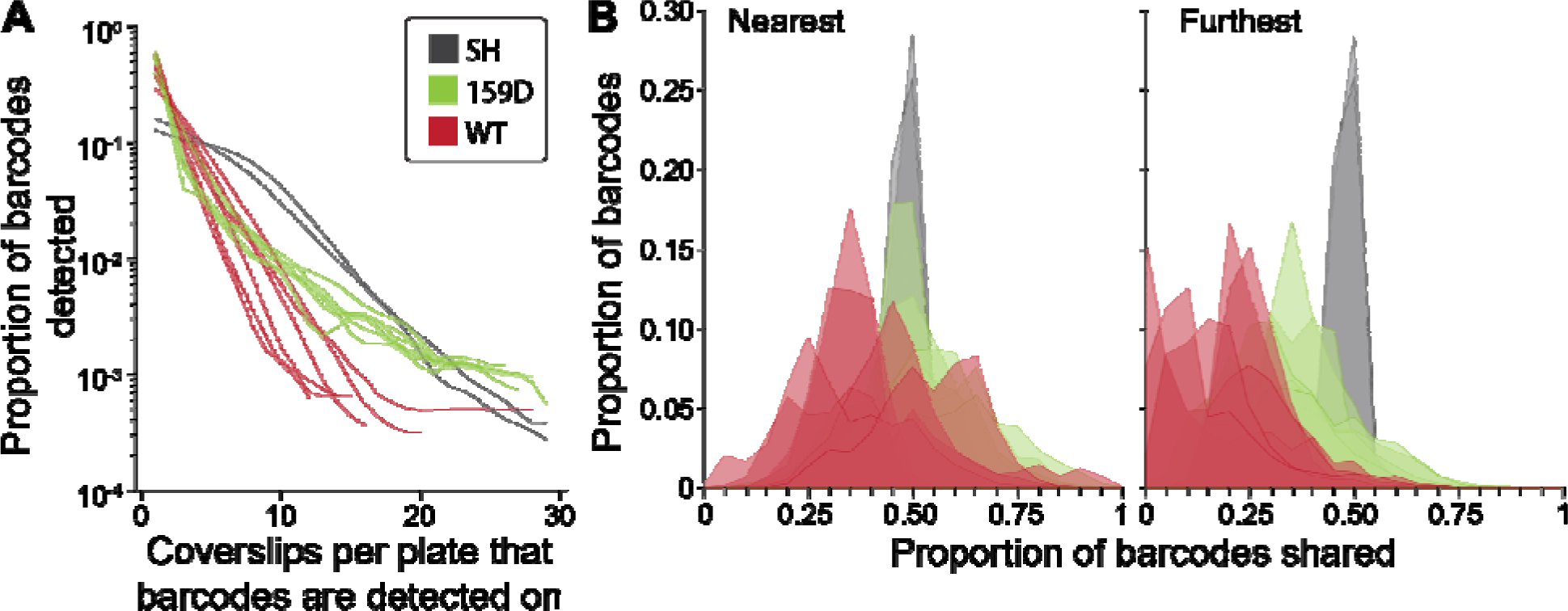
Viral phenotypic variation modulates the impact of spatial expansion on population genetic composition. A) The proportion of barcodes that are detected on a given coverslip is plotted against the number of coverslips per plate where that proportion is observed. Each line represents a single cell culture plate, with red and green colors representing different viral genotypes. Black lines show results from plates incubated infected with WT and incubated with constant shaking (SH), documenting results in the absence of spatial structure. Shading shows the 95% confidence interval of the distribution observed over 1000 sub-samplings of 3000 reads per coverslip. B) Histograms showing the proportion of barcodes shared between a given coverslip and the set of coverslips that are nearest (left) or furthest (right) from that coverslip. Each histogram represents a single plate. Colored histograms show results from different viral genotypes and black histograms show results obtained with WT virus under constant shaking.

## Discussion

We show that the spatial expansion of viral populations, even over a simplified landscape and involving only neutral diversity, is often accompanied by genetic bottlenecking and thereby gives rise to marked heterogeneity in population composition. Environmental constraints on dispersal augmented this effect, leading to greater dissimilarity between central and distal populations. In addition, changes to the viral dispersal phenotype, mediated through a substitution that is typical of seasonal antigenic evolution, modulated the genetic structure that developed through expansion. Given the central role of within-host expansion in acute viral infections, these results suggest that viral evolution may be shaped by stochastic processes accompanying dispersal.

Consistent with our observations, population genetic theory predicts that spatial expansion will lead to reductions in diversity due to the presence of small populations and therefore heightened genetic drift at the leading edge of the expansion (14, 28–30). This literature furthermore identifies several factors that are expected to modulate the extent to which diversity is lost during expansion. One key factor is the frequency and distance of dispersal events, described by the so-called dispersal kernel (31). The predicted relationship between genetic diversity and the number of long-distance dispersal events is non-monotonic, with an intermediate frequency of long-distance seeding events expected to precipitate the greatest reduction in population diversity (13). This effect follows from the rapid outgrowth of individual founders following dispersal to areas with abundant resources and no competition (13). Through the same mechanism, the dispersal kernel is also expected to shape the spatial heterogeneity, or patchiness, of an expanding population (32). Our data provide empirical support for these predictions, with dispersed viral subpopulations frequently showing marked reductions in barcode diversity relative to the source population, increased constraint on dispersal leading to more stringent bottlenecking, and a high degree of spatial heterogeneity forming across the landscape of expansion.

The population genetic structure that develops through influenza A virus dispersal *in vivo* has been examined in animal models (17, 19, 20, 33, 34). These studies monitored the spatial distribution of either multiple fluorescently tagged viruses in mice or a viral population carrying a genetic barcode in ferrets or guinea pigs. Relatively large spatial scales were evaluated with, for example, viral populations compared among five sections of the ferret trachea (20). At this spatial scale, genetic composition at different sites within the nasal cavity and the trachea is similar, revealing the maintenance of diversity during population expansion (17, 20). Sites within the lung show marked heterogeneity, however, both between and within individual lung lobes (17, 19, 20, 33, 34). Indeed, in ferret lungs, biopsies separated by ∼ 0.5–1 cm often showed highly distinct barcode compositions (17, 20).

These patterns observed *in vivo* show strong parallels with those seen in the cell culture model described here. The maintenance of diversity across well-separated locations in the upper respiratory tract and trachea is also seen in subsections of each culture plate. Both *in vivo* and in culture, these patterns indicate viral population expansion through processes of diffusive dispersal (20, 28). Conversely, the marked reduction in diversity that accompanies viral seeding of the lower respiratory tract suggests that long-distance dispersal is needed for the expanding viral population to occupy the lung lobes (20, 35). This process is mirrored in culture by the many subsections of the plates that show sharp reductions in barcode diversity. Both *in vivo* and in culture, these punctuated expansions yield isolated populations of distinct compositions. While these dynamics were observed using engineered diversity and experimental systems, analogous patterns in natural systems would give rise to within-host sub-populations of differing evolutionary potential. Indeed, spatial structure of within-host viral populations has been seen to shape pathogenesis (36), drug resistance (37) and variant transmission (38–40).

Within the respiratory tract, a complex array of biotic and abiotic factors is likely to drive viral dispersal. Biotic factors include the 3D structure of the infected tissue, the spatial distribution of susceptible cells, the viscosity and thickness of the mucus barrier (41), the presence of receptor decoys (22), and the directional flow of periciliary fluid driven by ciliary movement (25, 42). Many of these factors vary over the course of infection due to direct effects of the virus or indirect effects mediated by cytokine signaling (24, 43). Abiotic factors include Brownian motion, convection, shear forces mediated by airflow, and the behavior within the airways of respiratory particles produced by those shear forces (44). Only a subset of these factors is relevant within the cell culture model employed here. The absence of airflow and ciliary action are important to keep in mind: both are sources of directional movement, which may maintain population composition over larger distances. The lack of 3D structure and airway branching are also important considerations (45), with airway branching expected to impose genetic bottlenecking during viral dispersal (17). Given the simplicity of the cell culture model described, it is however well suited to the systematic introduction of new features and evaluation of their impact on population genetics.

Factors evaluated herein were the viscosity of extracellular fluids and the receptor binding properties of the virus. Both modulated population dynamics during expansion and we infer that each would also have an effect *in vivo*, increasing or decreasing the severity of within-host bottlenecks. Other factors that may change with viral strain or the intensity of host responses to infection include virion release via neuraminidase-mediated cleavage of receptors, the time needed for a newly seeded infection to spread (that is, the duration of the viral eclipse phase) and the timing with which infected cells become refractory to super-infection. While neuraminidase activity may modulate the relative frequency of diffusive and long-distance dispersal (46), eclipse phase duration and super-infection exclusion are expected to vary the opportunity for multiple long distance dispersal events to seed infection in the same area (13, 16).

In summary, our results align with the predicted consequences of spatial expansion for population genetic structure, revealing the random effects of genetic drift among other stochastic processes in defining the composition of dispersed subpopulations. Our data furthermore show that both viral and environmental features that modulate viral dispersal also modulate viral diversity and evolutionary potential, underlining the importance of biological context for defining the extent to which stochastic effects stemming from spatial structure are likely to shape viral evolution.

## Acknowledgements

We thank the staff of the Emory Integrated Genomics Core for their assistance with next generation sequencing and the staff of the Emory Integrated Cellular Imaging Core for their assistance with fluorescent imaging.

## Materials and methods

### Cells

Madin–Darby canine kidney (MDCK) cells were a gift from Dr. Daniel Perez, University of Georgia, Athens, GA. A seed stock of MDCK cells at passage 23 was amplified and maintained in Minimal Essential Medium (MEM; Gibco) supplemented with 10% fetal bovine serum (FBS; Atlanta Biologicals) and Normocin (Invivogen). 293T cells (ATCC, CRL-3216) were maintained in Dulbecco’s Minimal Essential Medium (Gibco) supplemented with 10% FBS and Normocin. All cells were cultured at 37 °C and 5% CO_2_ in a humidified incubator. The cell lines were not authenticated. All cell lines were tested monthly for *Mycoplasma* contamination while in use. The medium for the culture of influenza A virus in MDCK cells (virus medium) was prepared by supplementing Minimal Essential Medium with 4.3% bovine serum albumin (BSA; Sigma) and Normocin.

### Barcode

The construction of the plasmid library carrying the barcode, pDP Pan/99 NA-BC, was described previously (27). Briefly, the following DNA Ultramer (IDT) containing degenerate sites was synthesized: cacagtacatgataggaccccttaycgraccytattgatgaatgarttrggtgtyccattycayytrggracyaagcaagtgt gtatagcatggtcc. This product was then introduced into the cDNA form of the NA gene of influenza A/Panama/2007/99 virus by by Gibson Assembly (NEB).

### TC tag

A tetracystein tag was introduced into Pan/99 virus following the method of Han et al.(47). Briefly, in the context of the pDP Pan/99 NP plasmid, a flexible linker (GTGSGIR) followed by the 6 amino-acid Tc tag (CCPGCC) was inserted before the stop codon of the NP open-reading frame (GTGSGIRCCPGCC Stop). To maintain the NP vRNA packaging signal, the 200-nucleotide packaging sequence was duplicated (48). The construct was synthesized by Twist Bioscience.

### Viruses

Pan/99 NA-BC, Pan/99 NP-Tc and Pan/99 NA-BC HA-159D viruses were derived from influenza A/Panama/2007/99 (H3N2) virus and were generated by reverse genetics. In brief, 293T cells were transfected with eight reverse genetics plasmids. After a 16–24 h incubation at 37°C, the transfected 293T cells were co-cultured with MDCK cells in virus medium at 33°C for 40 h. Recovered virus was propagated in MDCK cells to generate working stocks. Propagation was carried out from low MOI to avoid accumulation of defective viral genomes but with a sufficient viral population size to maintain barcode diversity where present. Stocks were titered by plaque assay in MDCK cells. The presence of intended modifications and absence of *de novo* mutations was verified by sequencing of the viral genomes on an Illumina platform, following the MRT PCR method of Zhou and Wentworth (49). The presence of a diverse barcode in Pan/99 NA-BC and Pan/99 NA-BC HA-159D virus stocks was verified by focused next generation sequencing of the barcode region (see below).

### Growth kinetics

Replication of Pan/99 NA-BC and Pan/99 NA-BC HA-159D was evaluated in triplicate culture wells. MDCK cells in six-well dishes were inoculated at an MOI of 0.01 PFU/cell in PBS. After 1 h incubation at 33°C, inoculum was removed, cells were washed 3x with PBS, 2 mL virus medium was added to cells, and dishes were returned to 33°C. A 120 µL volume of culture medium was sampled at the indicated times points and stored at −80°C. Viral titers were determined by plaque assay on MDCK cells.

### Focus dispersion experiments

For experiments with Pan/99 NA-BC or Pan/99 NA-BC HA-159D viruses, 10 cm diameter round cell culture plates (Greiner) were prepared by using autoclaved centrifuge lubricant (ThermoScientific) to tack down sterile 8 mm glass coverslips (Harvard Apparatus). For experiments with Pan/99 NP-Tc, the same culture plates were used directly, without the placement of coverslips.

To prepare the center focus, MDCK cells (1 x 10^6^ in one well of a six-well dish) were inoculated with Pan/99 NP-Tc, Pan/99 NA-BC or Pan/99 NA-BC HA-159D virus at an MOI of 2 PFU/cell in 1x PBS (Gibco) and incubated at 33°C for 45 minutes. Warm virus medium was applied for 20 minutes before harvesting cells with 0.25% trypsin-EDTA (Gibco) and resuspending in growth medium. Cells were counted and brought to a concentration of 2.5 x 10^6^ cells/mL. Twenty microliters of cell suspension were pipetted in a droplet onto the coverslip tacked to the center of the plate, and 1 mL of 1x PBS was pipetted along the edge of the plate to prevent desiccation of the droplet. Plates were incubated at 37°C for 4 h to allow cells to adhere.

To complete the cell monolayer, after incubation, plates containing a center focus were washed once with 1x PBS before applying a 12 mL volume of MEM with 0%, 0.2%, 0.3%, or 0.4% v/v methyl cellulose and containing 5 x 10^6^ uninfected MDCK cells. Plates were incubated at 33°C in a standard CO_2_ incubator for 72 h, under static conditions or with constant agitation on a platform shaker. Agitation was achieved using a VWR Analog Rocking Platform shaker (VWR, 10127-872) with settings as 2 for speed and 8 for tilt which corresponds to 15 rpm and 12 degrees respectively.

For experiments with Pan/99 NA-BC or Pan/99 NA-BC HA-159D viruses, plates were then washed once with 1x PBS, and coverslips were collected. Cells were harvested from coverslips with 0.25% trypsin-EDTA, then the suspended cells were transferred to a 1 mL square-well 96-well block (Takara) used for RNA extraction. Blocks were centrifuged for 5 min at 1000 rpm to pellet cells, then supernatant was aspirated, leaving 140 µL of liquid per well for RNA extraction (see below). For experiments with Pan/99 NP-Tc, plates were instead processed by FlAsH staining as described below.

### FlAsH staining and fluorescent imaging

At the end of the 72 h incubation period, plates with Pan/99 NP-Tc virus were washed thrice with 6 mL 1x PBS, then stained with 2 µM FlaSH-EDT_2_ reagent (Cayman Chemical) diluted in Opti-MEM (Gibco, 3 mL solution per plate). During staining, plates were incubated at 37°C for 30 min. To remove FlaSH reagent, cells were washed once with 1x PBS, then thrice with 3 mL of BAL wash buffer diluted to 250 µM in 1x PBS and with incubation at 37 °C for 10 min per wash. Finally, cells were washed once with 1 mL of PBS, then fixed by addition of 3 mL 4% paraformaldehyde and incubation for 15 min at room temperature. After fixation, cells were washed twice with 1x PBS before visualizing using a BioTek Lionheart FX.

### Sample processing for next-generation sequencing of barcode region

The following method was used prepare samples for targeted sequencing of the barcode region of Pan/99 NA-BC. Viral RNA extraction (MN NucleoMag RNA extraction kit, Takara) was performed using samples transferred to 96-well blocks described under “Focus dispersion experiments,” followed by reverse transcription (Maxima RT, Thermo Fisher) with Oligo (dT) primer (49, 50). Viral cDNA was subjected to PCR amplification with primers flanking the barcode region. Samples were sent for amplicon sequencing to the Emory Integrated Genomics Core (EIGC). PCR to amplify the region containing the barcode was performed using PFU Turbo AD (ThermoFisher) with primers with partial sequencing adapters that generated a 155 nt product (P99_NA_adptr_428-449_F 5’-acactctttccctacacgacgctcttccgatctggaacaacactaaacaacaggc-3’ and P99_NA_adptr_563-582_F 5’-gactggagttcagacgtgtgctcttccgatctgcttttccatcgtgacaact-3’). Thirty PCR cycles were performed for all samples. Bead-based PCR purification was performed on all samples followed by quantification of DNA using Qubit. Samples were normalized to 20 ng/uL in nuclease-free water.

### Quantifying viral load by qPCR

Viral cDNA produced by RT (see above) was quantified by qPCR using SsoFast Evagreen Supermix (BioRad) according to the manufacturer’s instructions. Two technical replicates were performed for each sample and then averaged. Primers for Pan/99 NP were used (P99_NP_130F 5’-GTCGGGAAGATGATTGATGG-3’ and P99_NP_329R 5’-GGCCCTCCAGTTTTCTTAGG-3’).

### Analysis of barcode sequencing data

Sequences were processed using our custom software, BarcodeID, available in the GitHub repository: https://github.com/Lowen-Lab/BarcodeID and previously described (27). Briefly, BarcodeID uses BBTools to process raw reads, then uses a custom Python script to screen and identify barcode sequences present in each sample, then performs an error correction step to remove spurious barcode sequences, before finally calculating diversity statistics and writes summary tables. BBMerge screens for adapter sequences and merges forward and reverse reads using default settings, and BBDuk merges reads with a low average quality (<30). BarcodeID then screens each read to verify that the nucleotides at barcode and non-barcode sites match the nucleotides expected at those sites. Reads with mismatches are excluded from overall barcode sequence counts, but BarcodeID collects all high-quality variant amplicons and calculates overall mismatch rates by site to determine if any mutants with non-barcode alleles are driving any observed barcode dynamics. Samples with evidence of non-barcode driven dynamics were excluded from further analyses. BarcodeID contains an error correction step that has been previously described in detail (27).

## Data availability

All data generated or analyzed during this study are included in the manuscript. All raw sequencing data has been uploaded to NCBI’s Sequence Read Archive under the accession number PRJNA1529230. Original code developed in the course of this work can be found at https://github.com/Lowen-Lab/Spatial-barcodes_Holmes.

## Supplementary Figure Legends

**Supp Figure 1: Pan/99 NA-BC carries abundant synthetic diversity.** The barcode region lies within the NA gene segment and comprises 12 sites of synonymous variation. Two alleles are possible at each site, as shown schematically in the figure. Combinatorially, these polymorphisms yield a total of 2^12^=4096 unique haplotypes. The stacked bar plot shows the barcode composition of the virus stock used in experiments. Each color represents a distinct barcode, and its width shows its frequency in the population.

**Supp Figure 2: Comprehensive set of plots showing barcode composition and relative richness on cell culture plates (related to Fig 2 and 4).** Plates are grouped by condition, with the condition indicated at the upper left of each grouping. Within each grouping, plates are identified by a P number and paired pie chart and relative richness maps are arranged vertically. In the top row of each grouping, pie charts show barcode composition at 33 locations per cell culture plate. Each color represents a unique barcode and the size of the wedge indicates its frequency within the locally sampled population. In the bottom row of each grouping, a heat map shows barcode richness relative to the center focus, with circles scaled to viral load as measured with Ct values. P13, P05 and P06 are also shown in Figure 2.

