## Supplementary figures and images for "Viral dispersal drives changes in population composition"

### Supplemental Figure 1

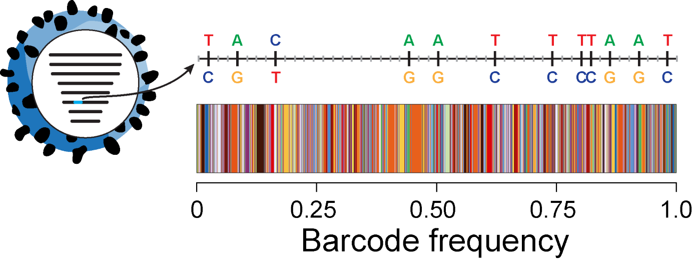

### Supplmental Figure 2

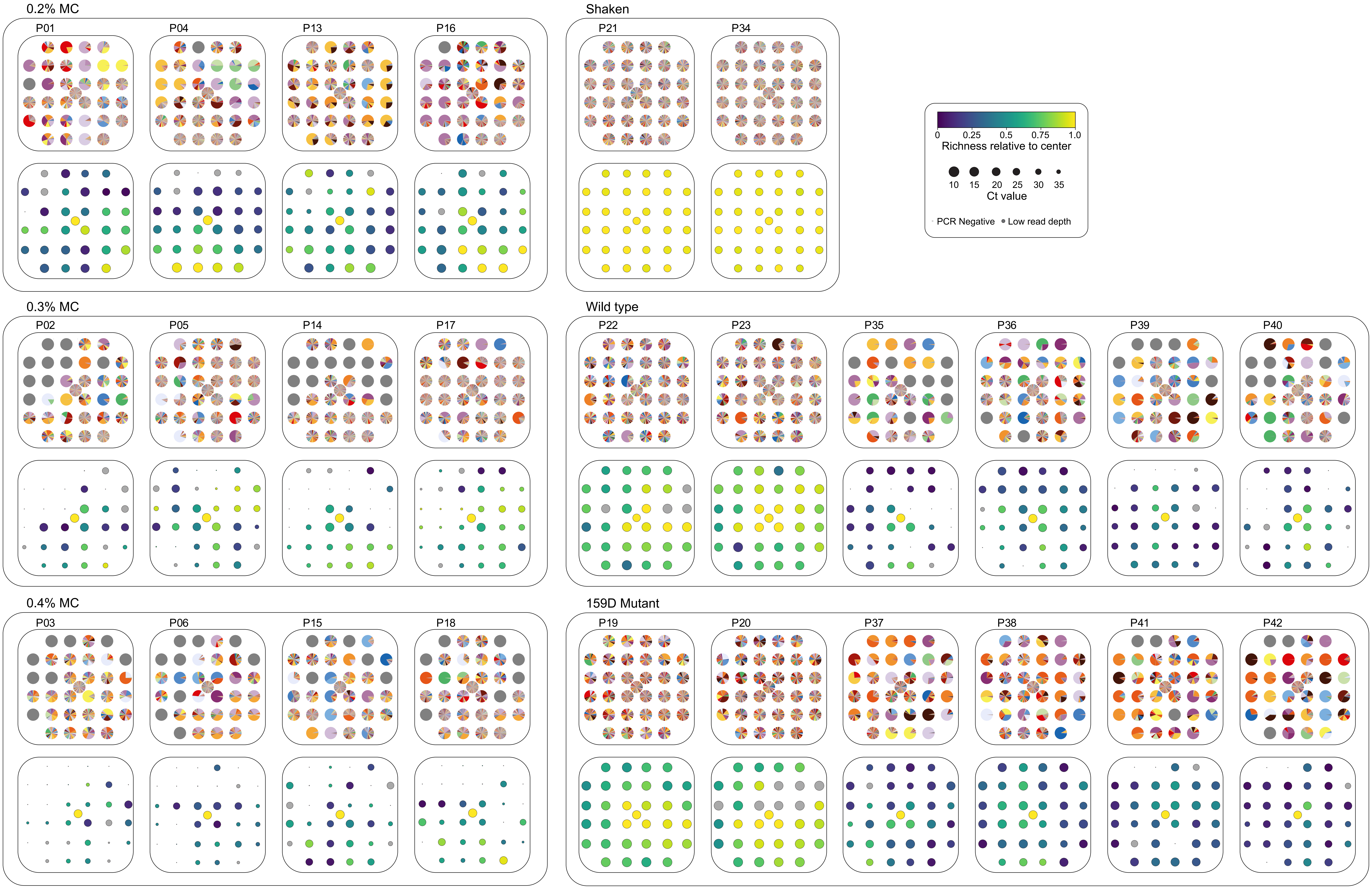
